# Electrophysiological signatures of spelling sensitivity development from primary school age to adulthood

**DOI:** 10.1101/2023.01.10.523398

**Authors:** Ekaterina Larionova, Anna Rebreikina, Olga Martynova

## Abstract

Recognizing spelling errors is important for correct writing and reading, and develops over an extended period. The neural bases of the development of orthographic sensitivity remain poorly understood. We investigated event-related potentials (ERPs) associated with spelling error recognition when performing the orthographic decision task with correctly spelled and misspelled words in children aged 8-10 years old, early adolescents aged 11-14 years old, and adults. Spelling processing in adults included an early stage associated with the initial recognition of conflict between orthography and phonology (reflected in the N400 time window) and a later stage (reflected in the P600 time window) related to re-checking the spelling. In children 8-10 years old, there were no differences in ERPs to correct and misspelled words; in addition, their behavioral scores were worse than those of early adolescents, implying that the ability to quickly recognize the correct spelling is just beginning to develop at this age. In early adolescents, spelling recognition was reflected only at the later stage, corresponding to the P600 component. At the behavioral level, they were worse than adults at recognizing misspelled words. Our data suggest that orthographic sensitivity can develop beyond 14 years.

## Introduction

One aspect of literacy is the ability to recognize errors in text. We check texts after writing or notice errors in words while reading. The search for mistakes in written words is closely related to the lexical representations of words stored in memory. Lexical representations of words can be divided into orthographic, phonological, and semantic components^1^. In addition to lexical processing, reading and spelling also involve sublexical processes responsible for spelling-to-sound mappings^2,3^. The event-related potential (ERP) method has good temporal resolution and makes it possible to evaluate various stages of processing verbal information at the millisecond level. When conducting ERP studies, multiple characteristics of words are manipulated to investigate the various temporal stages of processing visual-verbal information and the underlying cognitive processes. The sensitivity of ERP components to the phonological, lexical, and semantic characteristics of words varies at different stages of ontogenesis^4^. The process of recognizing the correct spelling can be associated with several stages of information processing – from sublexical processing to understanding the meaning of a word, which may differ in various age groups. However, this issue has not been sufficiently studied.

Many studies on the perception of orthographic violations examined evoked brain activity to pseudowords and non-words, i.e., letter strings that do not exist in the language. A word can turn into a non-word as a result of accidental typos. Previous ERP studies showed that brain responses to non-words differ from words at the sublexical processing stage by about 100-150 ms in fluently reading adults; the P1 component of ERPs was larger for atypical letter combinations than for typical ones^5–7^. Differences between words and non-words were also identified for components peaking at about 200 ms in adults (e.g., N170 or N1)^6,8^. Also, the N1 differed between words and pseudowords, which may be associated with the lack of visual representations for pseudowords in memory^5,7,9–11^. In adults, specific features in the processing of words, non-words, and pseudowords have also been found later, at the lexical-semantic processing stages corresponding to the N400 component^6,12–14^.

Sensitivity to orthographic structure in children develops in the early school grades as familiarity with letter sequences and with words and their spelling increases. In behavioral studies, words, pseudowords, and non-words can be reliably distinguished by the fourth grade (at the age of 9-11 years), when the reading of frequent words is already automated^15–18^. Remarkably, the word superiority effect in children of this age is less than in adults^19^. However, at the neural level, not much is known about the development of orthographic processing. Using the ERP in children aged 6-9 years old, differences in the N170 component were revealed between words and strings of symbols similar to the letters of the alphabet, but consisting of completely new characters^10,20–23^, however, there were no early differences between words, pseudowords, and non-words^18,20^. In the 11-year-old group, differences were found for the P150 and N400 components between pseudowords and non-words, and differences in 7-year-old children were revealed only at a later stage of information processing: the N400 was larger for non-words than words^18^. According to the authors, these results reflect lexical mechanisms of word superiority effects in younger children but both sublexical and lexical mechanisms in older children^18^. ERP differences for words, non-words, and pseudowords were also observed in children of different ages at later time intervals of word processing. In children aged 8-9 years, the amplitude of the late positive complex was significantly reduced for pseudowords compared to words^21,24^. The same patterns for a similar P600 wave were also obtained in children aged 12–14^25^. Adolescents (ages 15.4–19.3) show ERP patterns similar to adults at up to 110 ms in distinguishing between words and pseudowords that are earlier than in children^26^.

Unlike non-words and pseudowords, pseudohomophones, i.e., words with orthographic violations but pronounced like an existing word, are words with commonly occurring spelling errors. Therefore, using such stimuli can shed light on the mechanisms underlying one of the aspects of literacy – recognition of the correct and incorrect spelling of words. In various languages, many misspelled words tend to be phonologically similar to correctly spelled words. For example, Russian is a transparent language, but each vowel phoneme [i, e, a, ɔ] in unstressed syllables can be translated into two graphemes resulting in spelling errors. Thus, the pronunciation of the correctly spelled word “волна” (“wave”) with an unstressed first syllable, and the misspelled word “вална” (an incorrect spelling of a word “волна”) is the same [vɐɫˈna]. Similar phenomena can be observed in other languages. For example, in Spanish, there are several phonemes that can be translated into two or more graphemes (e.g., phoneme /b/ can be translated into b and v); these conflicting phonemes would produce an error in spelling^27,28^. For example, Spanish “baca” vs. “vaca” (“cow”) may be a frequent spelling error.

However, data from neurophysiological studies on the ERP components associated with the processing of pseudohomophones in adults are still inconsistent, and there are few ERP data are available for children. In adults, some studies have identified early perceptual effects of pseudohomophones at up to 200-250 ms^5,11,29–31^, but also, they were detected later, for the N400 and P600 components^21,22,30–32^. In the work of González-Garrido with colleagues, only late adolescents (16-18 years) with high performance on a 5-test battery of orthographic knowledge showed significant differences in P450 amplitude between correctly and incorrectly written words during the performance of an orthographic decision task. In contrast, adolescents with lower levels of orthographic knowledge showed no difference between the conditions^33^. There is evidence that the P600 component is higher for words and pseudohomophones than for pseudowords in 8-year-old children^24^. In 9–10-year-old children, differences between words and pseudohomophones have been observed: the amplitude of the late positive component was higher for correct words than for pseudohomophones^22^. Remarkably in 9-year-old children, the amplitude of the N400 component was higher for pseudowords than for words and pseudohomophones, but this effect was absent in children with spelling disorders^21^. Heldmann and co-authors^34^ studied the development of spelling sensitivity in elementary school in children of the 2nd (6.8-8.7 years) and 4th grades (9.6-10.8 years) during presentation of a line drawing (black lines on white background) representing a common object or an animal along with its proper correct or misspelled but phonologically correct name (e.g., a picture of a donkey with the correct German word for donkey “Esel”, or with a misspelled but phonologically correct version “Ehsel”) and found similar results to Bakos and colleagues^22^. The difference between correct and misspelled words was reflected by a more pronounced positive shift to misspelled words at about 600 ms, which was observed only in children in the 4th grade. However, differences in ERP between correct and misspelled words were also identified for 2nd graders but were observed at later stages, between 1300-1400 ms. The authors consider this effect a marker of orthographic sensitivity and conclude that there is a distinct shift from the 2nd (age range 6.8-8.7) to the 4th (age range 9.6-10.8) grade with regard to orthographic sensitivity^34^. Gómez-Velasquez and colleagues examined the detection of spelling errors in 8-year-old children with varying performance on four naming tasks (drawings, letters, numbers, and colors)^35^, and their results partially agree with the results of Heldmann and colleagues^34^. Their study found enhanced amplitude for negativity peaking at 380 ms (N380) and also enhancement of the subsequent positive component (600-700 ms) for pseudohomophones in children with average naming performance that was not found in children with slow naming performance^35^. Thus, ERPs in children of the 2nd grade with good reading skills in Gomes-Velaskes’s study^35^ were similar to ERPs in children of the 4th grade in Heldmann’s study^34^. It is important to note that in the experiment of Gomes-Velaskes^35^ and in the Heldmann study^34^, the presentation of verbal stimuli was associated with the presentation of the corresponding picture. This means that such paradigms differ from those used in studies with adults, which makes it difficult to compare the results obtained in different age groups. At the same time, using corresponding pictures could facilitate the recognition of the correct spelling of words in children. As far as we know, a comparative study of orthographic sensitivity that includes both groups of children and adults has not been previously conducted. Therefore, one of the objectives of our study was to use the same paradigm to study different age groups and not to use corresponding pictures that could potentially lead to improved spelling recognition.

Another significant gap in the knowledge of the neural underpinnings of literacy is the lack of ERP data in processing pseudohomophones in adolescents. Most studies of the neurophysiology of reading in children reported ERP findings for children in grades 1-5, that is, during the reading development period^21,22,24,34–36^. The development of specialized processing of lexical units and the differentiation of words from non-words, pseudowords, and more complex units – pseudohomophones reflect the automation of reading processes^37,38^. Behavioral studies have shown that children in the 1st grade respond faster and more accurately to real words than simple pseudowords^39^. And children 8–10 years old are already able to distinguish between valid and invalid combinations of letters in pseudowords if they have the opportunity to choose between two alternatives (for example, sinnum or ssinum)^40^. But it is not until 4th grade that children use phonological and orthographic patterns to distinguish between pseudowords and letter strings in the same way adults do^39^. Thus, behavioral data show that automatic word recognition develops gradually, starting in early school age, and reaches the adult level, at least for simple words, by grade 4^17,36,41–43^. At the same time, there are neurophysiological data that demonstrate that 5th-grade children still differ in ERP word processing patterns from adults, and the path to fluent reading continues beyond 5th grade^18^. In addition, even in good readers aged 10-11 years the functional lateralization of linguistic neural networks involved in automatic word recognition and phonological processing is still not developed^44,45^. Although neurophysiological evidence suggests that early adolescents use similar strategies to adults in processing and learning new words and can effectively use context to anticipate incoming information^46,47^, the visual word processing system continues to develop^48^.

Moreover, ERP studies have shown non-linear complex dynamics in the development of processing of various characteristics (spelling, lexical, semantic) of verbal stimuli; for example, in a study by Coch and Benoit^36^, 4th-grade students showed smaller differences in ERPs between words, pseudowords, nonpronounceble letter strings, and false font strings than 3rd and 5th-grade students. Changes in the nature of errors in written language as children grow older are also noted: shifts in the proportion of spelling and morphological errors were observed between grades 4 and 5, and the relative frequency of morphological errors increased in older school students^49^. The authors hypothesized that the normal development of spelling reflects non-linear growth and that it takes a long time to develop a robust spelling vocabulary that coordinates phonology, spelling, and morphology and supports word-specific, regular spelling. Therefore, studies in various age groups are needed to form a complete picture of the age dynamics of the formation of literacy.

In this study, we investigated the process of spelling error recognition when presented with correctly spelled words and words with real spelling errors in the orthographic decision task and features of spelling recognition in different age groups. This study aimed to compare ERP patterns associated with spelling recognition in three age groups: primary (8–10 years old) and middle-school-aged children (early adolescence, 11–14 years old) and adult native speakers. An important feature of this study is the use of words with real misspellings rather than artificially constructed pseudo-homophones that do not occur in natural written speech. We assume that as the processes of spelling and reading are automated, earlier neurophysiological markers of recognition of spelling errors will be revealed. If the formation of spelling sensitivity is completed by early adolescence, then middle-school-aged children will have similar spelling recognition ERP patterns to adults.

## Methods

### Participants

We investigated three age groups: primary and middle-school-aged children (early adolescence) and adult native speakers. Twenty-seven healthy children aged 8 to 10 years old (20 female, 7 males; mean age 8.8, SD 0.9; educational level 2.8, SD 0.8 years), twenty-five healthy children aged 11 to 14 years old (9 female, 16 males; mean age 12.7, SD 0.9; educational level 6.6, SD 0.8 years) and thirty-six adult healthy volunteers aged 18 to 39 years old (22 female, 14 males; mean age 24.5, SD 5.0; educational level 13.9, SD 1.8 years) participated in the study. All participants were right-handed, native speakers of Russian, without speech disorders, neurological, or psychiatric diseases. They had normal or adjusted to normal vision. All children aged 11 to 14 years old, with the exception of one, had good academic performance in the Russian language (they had a score of 4-5 points, where 5 is the maximum score). Children aged 8-10 were not assessed in Russian language at school, so we relied on parental reports that children did not experience spelling difficulties. All participants or their legal representatives in the case of children gave written informed consent to participate in the study. The study was carried out in compliance with the Declaration of Helsinki and was approved by the Ethics Committee of the Institute of Higher Nervous Activity and Neurophysiology of the Russian Academy of Sciences (protocol #03 from 15.07.2019).

### Stimuli Material

The stimuli were 99 singular and plural nouns spelled correctly and incorrectly. Each misspelled word contained only one error. To neutralize the influence of the word length factor on processing, all selected words were 5–6 letters long. Two types of stimuli were presented: words with unstressed vowels in the word root written correctly (correct root [CR], 50 words; for example, “волна” [vɐɫˈna] [“wave”]) and incorrectly (misspelled root [MR], 49 words; for example, “блоха” [bɫɐˈxa] [“flea”]). An important feature of these stimuli is that the correctly spelled words and the misspelled words sound the same. For this reason, errors often occur in words with an unstressed vowel. A complete list of the stimuli and their characteristics is provided in Appendix A.

The frequency of words was determined using the Frequency Dictionary of the Modern Russian Language^50^. The mean frequency of CR words was 110.77 (range 0.7-926), while that of MR words was 81.98 (range 1.5-709). The CR and MR words were equalized in frequency (Т = 555.0, p = 0.57, level of significance: 0.05). We also compared the orthographic neighborhood size using the StimulStat database^51^; this parameter did not differ statistically for correct and misspelled conditions (T = 151.5, p = 0.54). The mean orthographic neighborhood size of CR words was 1.0 (SD 1.0), MR 1.0 (SD 1.0). The frequency of bigrams containing an error for misspelled words, or bigrams containing an unstressed vowel, which could have been misspelled for a correctly spelled word was equalized for correct and misspelled words (T = 579.0, p = 0.74): the mean bigram frequency of CR words was 3061677 (SD 1554942), MR 2975152 (SD 1574369) (determined according to Frequency Dictionary of Modern Russian Language^50^).

### Experimental Procedure

During the EEG recording, the participants sat in a comfortable chair in a darkened room at a distance of about 1 m from the computer screen. The subjects were instructed to silently read the words presented on the screen and determine whether the word on the screen was spelled correctly or misspelled by pressing the left or right buttons of a Logitech F310 gamepad. The allocation of right and left buttons for correct and misspelled words were balanced across subjects.

The words were presented in the center of the screen in a white font (lower case Liberation Sans (an analog of Arial), 125 pt) on a black background. Stimuli were presented in a random order on a 19 “ LG FLATRON L1952T monitor using PsychoPy Experiment Builder v3.0.7 software^52^. Stimulus presentation began with a fixation cross; stimuli were presented on the screen until the subject responded, the time interval after the response until the next stimulus varied from 1300 to 2300 ms. The total time for completing the task did not exceed 40 minutes for children aged 8 to 10 years old, 15 minutes for children aged 11 to 14 years old and 10 minutes for adults.

### EEG Recording and Data Processing

EEG was recorded from 19 electrodes Fp1, Fp2, F3, F4, F7, F8, C3, C4, T3, T4, T5, T6, P3, P4, O1, O2, Fz, Cz, Pz, according to the International 10–20 system guidelines, referenced to the mastoids. Data were sampled at 250 Hz; electrode impedances were below 10 kΩ.

Offline processing was carried out using Brain Vision Analyzer 2.0.4 software (Brain Products, GmbH, Munich, Germany). Offline filtering was performed using a band-pass filter (0.5–30 Hz), followed by fast ICA for removing eye-blink artifacts. Data were segmented into epochs starting 300 ms before the word onset and lasting until 1500 ms after the onset. Only correct trials were segmented. Semi-automatic artifact rejection (+/−100 μV threshold) was performed on each epoch section (−300–1500 ms). Data were corrected relative to a 300 ms prestimulus baseline.

Data were averaged per subject and per condition. Five children aged 8 to 10 years were excluded due to an insufficient number of averagings (less than 28 realizations) and excessive noise or artifacts. Three adult participants were excluded due to excessive noise or artifacts. In section 2.1, data are provided only for those volunteers whose data were ultimately analyzed.

The reaction time and error rate were analyzed using ANOVAs with repeated measures (RM) for three groups of participants. One participant whose average reaction time exceeded three standard deviations (from the group of children aged 8-10 years old) was excluded from the analysis of behavioral data. The reaction time was only evaluated for correct answers. All significant (p < 0.05) main and interaction effects were followed by post hoc Bonferroni-corrected contrasts. To correct violations of sphericity and homogeneity, the Greenhouse–Geisser correction was applied as well. Statistical analysis was performed using the STATISTICA software (Statsoft, Tulsa, OK, USA).

Statistical analyses of the ERPs were carried out using the Matlab FieldTrip toolbox. A permutation t-test was performed to explore significant differences between CR and MR conditions in the window between 0 and 900 ms after the stimulus for the adult group and in the window between 0 and 1500 ms after the stimulus for the two groups of children, whose response time was longer compared to adults (Fieldtrip, Monte Carlo method, 500 permutations at 19 electrodes). The differences between CR and MR words were considered significant if maintained for a minimum of 5 consecutive samples (i.e., over 20 ms) in at least 2 neighboring electrodes and with an alpha level of 0.025.

## Results

### Behavioral Data

The RM ANOVA results are shown in Figure 1. The analysis of error rate indicated a significant main effect for Group F(2,84) = 28.46, р < 0.0001, η2p = 0.40: the total percentage of errors for CR and MR differed in all three groups (adults vs. early adolescents vs. children, 2.65 vs. 8.68 vs. 13.91 %). A significant Spelling × Group interaction was also found F(2,84) = 10.69, р < 0.0001, η2p = 0.20: the percentage of errors for MR differed between the adult and early adolescent groups (adults vs. early adolescents, 3.97 vs. 13.68 %), the percentage of errors for both CR and MR differed between the adult and children groups (CR: adults vs. children, 1.33 vs. 9.54 %; MR: adults vs. children, 3.97 vs. 18.29 %), the percentage of errors for CR differed between the early adolescent and children groups (early adolescents vs. children, 3.68 vs. 9.54 %).

**Figure 1.**
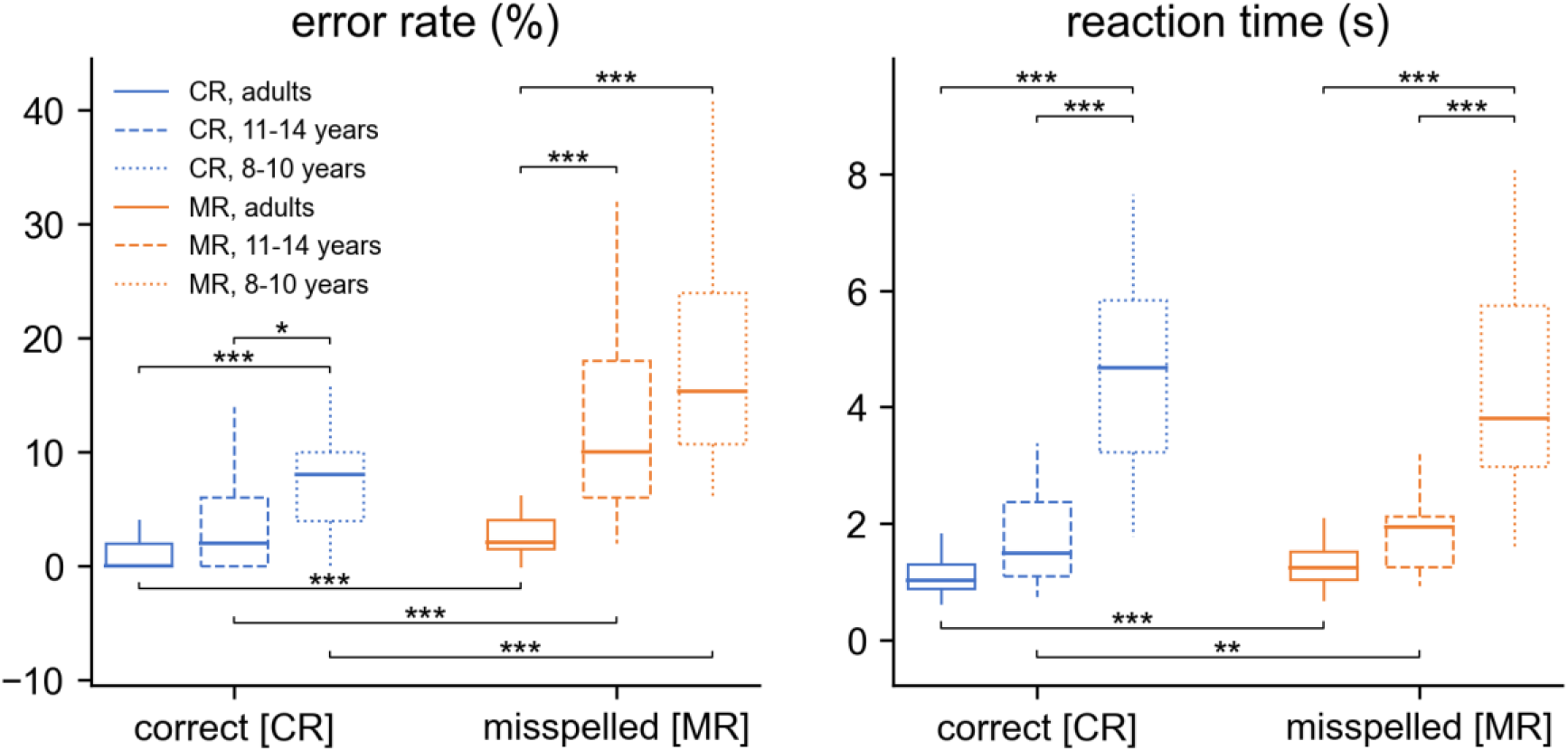
RM ANOVA results and post-hoc t-tests within each group for error rate and response time. Note: * denotes p < 0.05, ** denotes p < 0.01, *** denotes p < 0.001.

Dependent post-hoc t-tests within each group revealed that adults, early adolescents and children gave more correct answers to CR than MR (adults: CR vs. MR, 1.33 vs. 3.97 %; early adolescents: CR vs. MR, 3.68 vs. 13.68 %; children: CR vs. MR, 9.54 vs. 18.29 %).

The analysis of reaction time indicated a significant main effect for Group F(2,84) = 71.70, р < 0.0001, η2p = 0.63: the total reaction time for CR and MR differed between adults and children (1.17 vs. 4.55 s), and between early adolescents and children (1.80 vs. 4.55 s). Significant Spelling × Group interaction was also found F(2,84) = 7.24, р < 0.001, η2p = 0.15: the reaction time for both CR and MR differed between adult and children groups (CR: adults vs. children, 1.09 vs. 4.68 s; MR: adults vs. children, 1.25 vs. 4.41 s), the reaction time for both CR and MR differed between the early adolescent and children groups (CR: early adolescents vs. children, 1.70 vs. 4.68 s; MR: early adolescents vs. children, 1.89 vs. 4.41 s).

Post-hoc t-tests within each group revealed that only adults and early adolescents had longer reaction times for MR than CR (adults: CR vs. MR, 1.09 vs. 1.25 s; early adolescents: CR vs. MR, 1.70 vs. 1.89 s).

### ERP analysis

We found differences between ERPs to CR and MR conditions in adult participants in two time windows (Figure 2). More negative activity distributed across the frontal and posterior scalp sites was found for misspelled than for correctly spelled words around 400 ms that is compatible with N400 (300-520 ms, p = 0.002). More positive activity distributed across the frontal, central and posterior scalp sites was found for misspelled words than for correctly spelled words around 700 ms that is compatible with P600, (592-868 ms, p = 0.002).

**Figure 2.**
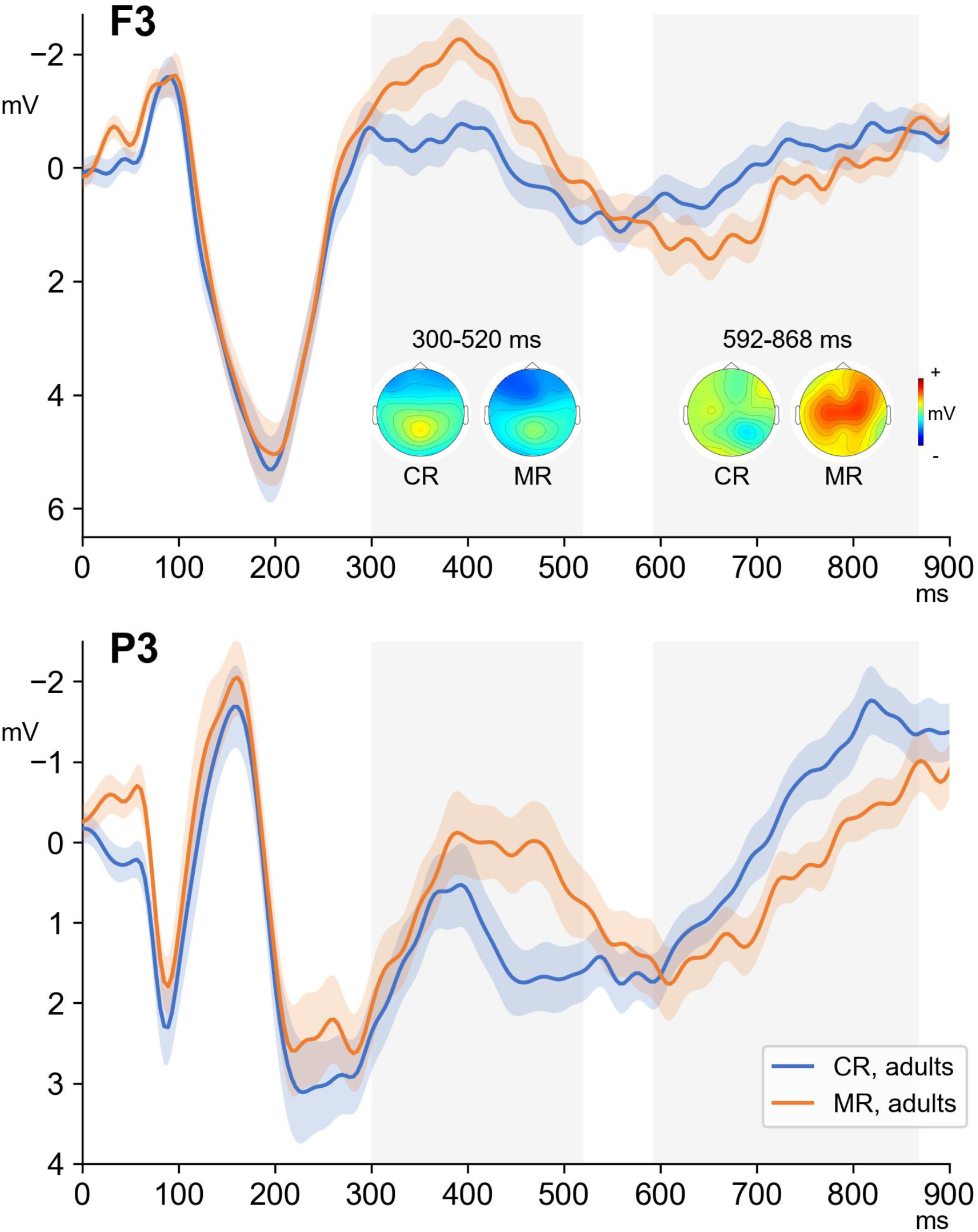
Averaged ERPs and topographic maps for correct (CR) and misspelled (MR) words in adult subjects for P3 and F3.

We found differences between ERPs to CR and MR conditions in children aged 11 to 14 years in the period from 768 to 1192 ms (p = 0.002). In this group, more positive activity, predominantly distributed across the central scalp sites was found for misspelled words than for correctly spelled words that is compatible with P600, (Figure 3). However, we did not find any difference between ERPs to CR and MR conditions in children aged 8 to 10 years (Figure 3).

**Figure 3.**
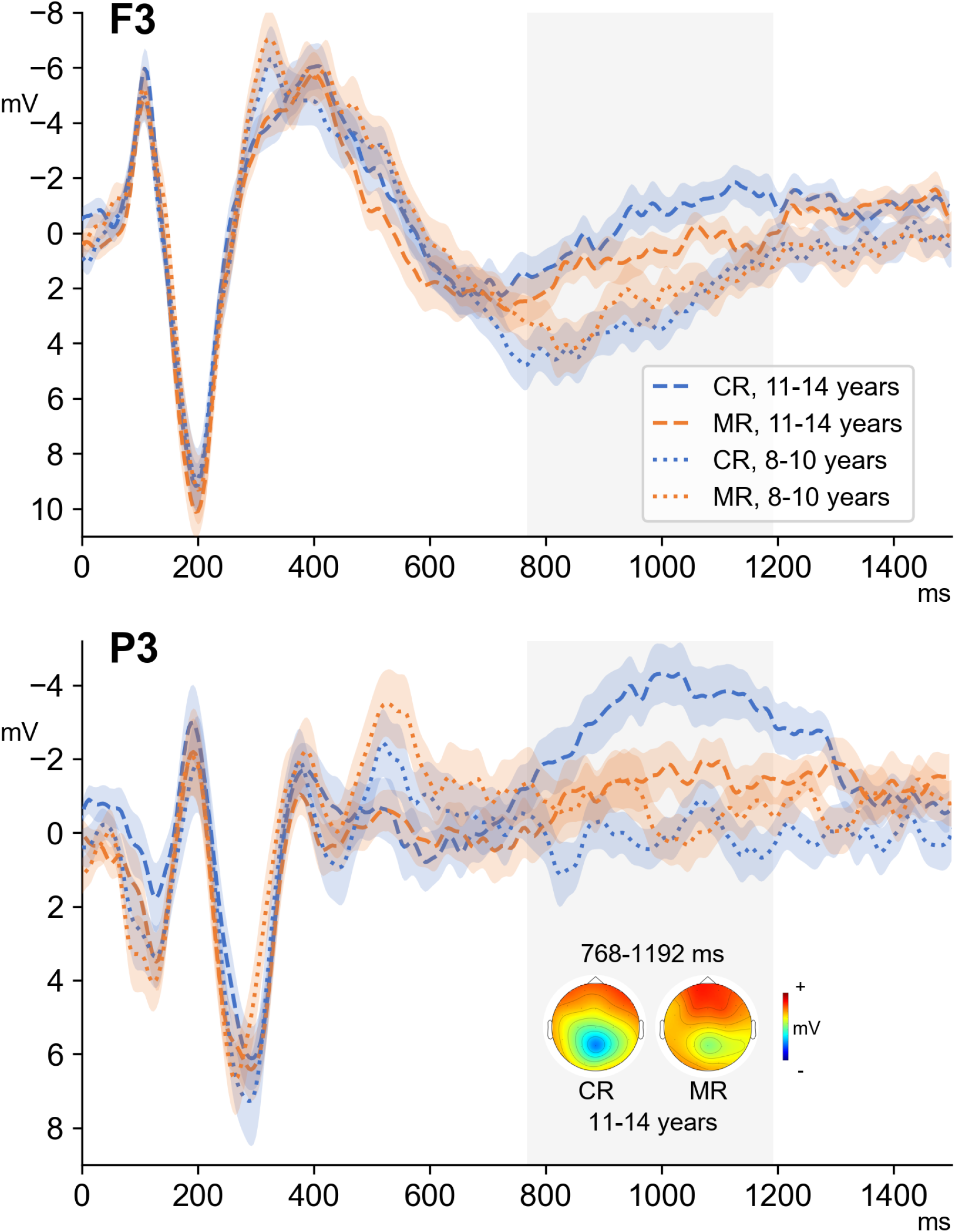
Averaged ERPs and topographic maps for correct (CR) and misspelled (MR) words in children for P3 and F3.

## Discussion

We investigated the neural processes that underlie recognizing the correct spelling of words and the features of spelling processing in different age periods (adults, 11-14 years old children, and 8-10 years old children). At the behavioral level, all the studied groups successfully identified correctly spelled words and misspelled words, the average percentage of errors even in the group of children aged 8-10 did not exceed 14%. Children had more incorrect answers and longer reaction times than early adolescents and adults. Early adolescents and adults showed similar, but not identical, behavioral results. The response time to both types of stimuli and the percentage of erroneous answers to correctly spelled words did not differ between these groups, at the same time early adolescents were worse than adults at recognizing misspelled words. At the ERP level, it has been shown that adults demonstrate earlier spelling error recognition patterns compared with early adolescents 11-14 years old, corresponding to the N400 (300-520 ms) and P600 (592-868 ms) components. In the early adolescents, the recognition of spelling errors was associated only with the late positive wave (768-1192 ms). In children of 8-10 years old, no differences were found in ERPs for correctly and incorrectly spelled words. Thus, our results show that orthographic sensitivity can develop until at least 14 years of age.

A higher percentage of errors was observed for misspelled words compared to correct ones, which is consistent with many previous studies^9,53–55^. It has previously been shown that correctly spelled words are recognized better and faster than misspelled words in adult native speakers and in late adolescents^31,33,56^. It is important to note that in our study adults and early adolescents did not differ in response time to either type of stimuli: they equally quickly recognized correctly spelled and misspelled words, that is, they probably used similar reading and spelling recognition strategies. However, the percentage of misidentified misspelled words was higher in early adolescents than adults, which may indicate that spelling sensitivity is still developing at this age.

Interestingly, in our work, the reaction time for misspelled words and correctly spelled words differed only in the groups of adults and early adolescents but not in children aged 8–10 years. The dual-route model of reading supports either an indirect phonological route or a direct orthographic route for semantic access^2,57^. Misspelled words do not have orthographic representations and are therefore likely to activate an indirect route, which slows down recognition, while correctly spelled words activate a direct route in experienced readers^58^, which is consistent with our findings in adults and early adolescents. At the same time, phonological skills are primary in reading acquisition when an orthographic representation is not fully specified, and novice readers predominantly use an indirect phonological route for lexical access to correctly spelled words^59–61^. Behavioral evidence demonstrates that 3rd grade readers typically rely predominantly on a direct lexical strategy to read familiar words silently, unlike grade 2 children^62–64^. In addition, children at the end of grade 3 (mean age 9.5) may already show faster reaction times for words compared to pseudo-homophones in the phonological decision task^22^. Thus, in our study, the sample of children aged 8–10 years (mainly pupils in grades 2 and 3) corresponds to the period of transition from the indirect phonological route to the direct orthographic route for semantic access, and the obtained behavioral results may reflect the dominance of the phonological route in children aged 8–10 years. Our behavioral data is broadly consistent with the electrophysiological evidence we received.

For the N400 component, we found differences between correctly spelled and misspelled words only in adults: the N400 amplitude across the frontal and posterior scalp sites was more negative for misspelled words. The N400 component is associated primarily with lexical-semantic information processing and reflects the integration of orthographic and phonological information with lexico-semantic representations^6,12–14^. Furthermore, the frontal N400 is associated with familiarity memory: its amplitude decreases for familiar stimuli^65,66^. Pseudohomophones evoked an increase in the amplitude of N400 compared to words in adult native speakers and late adolescents^30–32,67^ that is probably associated with lexical-semantic conflict and difficulty in semantic access. Gonzalez-Garrido and colleagues^30^, suggested that recognizing an error causes a conflict, which can be resolved by visually re-comparing the word with its stored visual representation. So, the enhanced N400 for misspelled words may reflect conflict in the integration of the visual and phonological forms and meaning-based representations. On the other hand, N400 (FN400) has been associated with familiarity memory, and even implicit memory^68–70^. So our findings may also reflect inconsistency in incoming visual information and visual word forms stored in memory.

The revealed absence of the N400 effect in early adolescents may indicate that orthographic representations for correctly spelled words are not strong enough therefore there is no conflict between orthographic and phonological word form or between sensory input and memory when processing misspelled words. As there is some evidence that FN400 can be related to implicit memory, we hypothesized that in adults the more prominent N400 for misspelled word may reflect an implicit process, while recognizing spelling in early adolescents is less automated and related only to explicit analysis in the time window of 768-1192 ms, corresponding to the P600 wave. Nonetheless, the process of spelling recognition in adults is not limited to the time window of 300-520 ms; they demonstrate neural patterns similar to early adolescents for the late wave corresponding to the P600 component.

The functional role of the P600 component is still a matter of debate. P600 refers to access to the phonological lexicon and knowledge of word spelling^24^. Moreover, P600 reflects re-processing to check for possible processing errors^71,72^. González-Garrido and colleagues^30^ suggested that P600 could be related to parsing, which finally made it possible to distinguish between words and pseudo homophones. Several studies have found that recollection is associated with a 500–700 ms parietal effect termed the late-positive complex (LPC or P600)^66,73^. Recollection involves the explicit retrieval of specific details about something recognized^74^. Consequently, in our study, the incomplete coincidence of “memory traces” for the visual word forms and the later explicit “rechecking” of the presence of errors (or recollection of details of word spelling) can explain the larger P600 amplitude for misspelled words in adults. We assume that the P600 component in early adolescents has a different functional meaning than P600 in adults and is not associated with re-processing to check for possible processing errors, but reflects only primary explicit analysis (without rechecking), so at the behavioral level, early adolescents recognize misspelled words worse than adults do.

Thus, the adult spelling recognition process involves at least two steps: implicit detection of conflict between orthography and phonology associated with the N400 component and subsequent explicit rechecking for errors related to the late positive component. In the early adolescence, recognizing misspellings is associated only with explicit analysis of words reflected in the late positive wave. Somewhat consistent data were obtained by González-Garrido and colleagues: the N400 component differed between correctly and incorrectly spelled words only in adults with good spelling skills, while the P600 component differed between correctly and incorrectly spelled words in both good and poor spellers^30^. In our study, early adolescents, who are less experienced in reading and spelling, had differences between responses to MR and CR in the P600 component, and more experienced adult readers had differences in both N400 and P600.

Children 8-10 years old demonstrate similar neural processes for correctly spelled words and words spelled incorrectly, at least up to 1500 ms, as well as a longer process of recognizing spelling and finding the correct answer at the behavioral level. Comparable reading processes can explain this for correctly spelled and misspelled words, while it is likely that children need more time to determine the correct spelling (longer than the 1500 ms epoch we analyzed), which is consistent with the long response time to both types of stimuli. Similar results were found in the phonological decision task: neither N400 nor LPC (similar to P600 wave in our study) differed between words and pseudohomophones in children of 8 years old (although at the behavioral level children recognized words better than pseudohomophones)^24^. However, in the other study, older children aged 9-10 in the phonological decision task already showed higher average LPC amplitudes (600-976 ms) for words compared to pseudohomophones^22^. It could be assumed that the spelling task, unlike the phonological task, modulates the differences between words and pseudo-homophones, but our data showed that the spelling task did not modulate the differences in ERP in children. At the same time, as we already noted in the introduction, in the few studies with the spelling task in which differences between words and pseudohomophones were observed, images were used that were presented before a verbal stimulus and probably, facilitated the detection of conflict between orthography and phonology in children 8-10 years old^34,35^. The division of children aged 8–10 years into narrower age groups may be useful in further research since, during this period, the initial patterns of recognition of correct spelling at the neural level might be formed. However, dividing our sample into subgroups of children aged 8, 9, and 10 did not show significant differences in ERPs between correctly spelled and misspelled words (see Appendix B). The absence of differences could be due to the small number of children in the subgroups.

## Conclusion

The present ERP study provides electrophysiological data showing age-related differences in spelling error recognition. In adult native speakers, ERP analysis of brain responses to correctly spelled and misspelled words revealed two stages of error processing: an earlier one associated with the initial recognition of conflict between orthography and phonology (reflected in the N400 time window) and a later one (reflected in the P600 time window), probably related to re-checking for errors in the spelling of the word. In early adolescents aged 11-14 years, spelling recognition is reflected only at a late stage, corresponding to the P600 component. We did not find differences in the ERP in the time window up to 1500 ms in children aged 8-10 years old. Our results imply that orthographic sensitivity begins to develop in primary school and does not allow preattentive recognition of the correct spelling (without attentive rechecking of the correctness of spelling). Thus, our results confirm previous adult findings and provide new data on the neurophysiological underpinnings of spelling processing in children and adolescents. It is important to note that we have shown that orthographic sensitivity can develop until at least 14 years of age. However, further research is needed to determine at what stage the formation of orthographic sensitivity is completed, and the transition to at least two-stage recognition of the correctness of spelling, similar to adult native speakers, is realized.

## Data availability

The datasets generated during and/or analyzed during the current study are available from the corresponding author on reasonable request.

## Acknowledgements

Grateful thanks are due to the parents and children who elected to participate in this study. This study was partially supported by grant No. 20-013-00514 of the Russian Foundation of Basic Research (RFBR) and IHNA & NPh RAS.

## Contributions

E.L., A.R. and O.M. conceived the experiment, E.L. and A.R. conducted the experiment and analyzed the results. E.L. produced the first draft of the manuscript and all authors contributed to writing and revising the paper. All authors reviewed the manuscript before submission.

## Ethics declarations

Competing interests

The authors declare no competing interests.

## Supplementary Information

Supplementary Information 1. Stimulus list with relevant characteristics.

Supplementary Information 2. The averaged ERPs for correct (CR) and misspelling (MR) words in children aged 8, 9, and 10.

