## Supplementary Information 1. for "Electrophysiological signatures of spelling sensitivity development from primary school age to adulthood"

Stimulus list with relevant characteristics.

Note: Type – type of the stimulus; CF – correct form for MR; GN – grammatical number; Fr – frequency, ipm – instances per million words; L – length; NS – number of syllables; IPA – The International Phonetic Alphabet; EP – error position for MR

| **Word** | **Type** | **CF** | **GN** | **Fr, ipm** | **L** | **NS** | **IPA** | **EP** | **Word sense** |
| --- | --- | --- | --- | --- | --- | --- | --- | --- | --- |
| бл**и**ны | CR |  | plural | 12.8 | 5 | 2 | [blʲɪˈnɨ] |  | pancakes, thin batter cakes |
| б**о**ксёр | CR |  | singular | 5.3 | 6 | 2 | [bɐˈksʲɵr] |  | boxer |
| в**е**сна | CR |  | singular | 91.3 | 5 | 2 | [vʲɪˈsna] |  | spring (season) |
| в**и**нты | CR |  | plural | 9.4 | 5 | 2 | [vʲɪnˈtɨ] |  | screws, propellers |
| гл**а**за | CR |  | plural | 857 | 5 | 2 | [ɡɫɐˈza] |  | eyes |
| гл**о**ток | CR |  | singular | 15.8 | 6 | 2 | [ɡɫɐˈtok] |  | swallow, draught, gulp |
| б**о**рьба | CR |  | singular | 190.5 | 6 | 2 | [bɐrʲˈba] |  | struggle, fight, conflict, combat |
| бл**о**ха | CR |  | singular | 4.3 | 5 | 2 | [bɫɐˈxa] |  | flea |
| гр**и**бы | CR |  | plural | 38 | 5 | 2 | [ɡrʲɪˈbɨ] |  | mushrooms |
| д**и**карь | CR |  | singular | 7.4 | 6 | 2 | [dʲɪˈkarʲ] |  | savage, shy person, unsociable person |
| бр**е**вно | CR |  | singular | 22.5 | 6 | 2 | [brʲɪˈvno] |  | log, beam (trunk of dead tree, cleared of branches) |
| д**о**ска | CR |  | singular | 67 | 5 | 2 | [dɐˈska] |  | board, plank, blackboard |
| др**о**ва | CR |  | plural | 25.8 | 5 | 2 | [drɐˈva] |  | firewood |
| ж**е**лток | CR |  | singular | 3.3 | 6 | 2 | [ʐɨɫˈtok] |  | yolk, vitellus |
| в**е**дро | CR |  | singular | 34 | 5 | 2 | [vʲɪˈdro] |  | bucket, pail |
| з**е**лень | CR |  | singular | 24.6 | 6 | 2 | [ˈzʲelʲɪnʲ] |  | verdure (greenness, vegetation) |
| к**а**ток | CR |  | singular | 7.8 | 5 | 2 | [kɐˈtok] |  | skating rink, ice rink |
| вол**о**сы | CR |  | plural | 141 | 6 | 3 | [ˈvoɫəsɨ] |  | hair |
| кр**е**пёж | CR |  | singular | 0.7 | 6 | 2 | [krʲɪˈpʲɵʂ] |  | fastening, strengthening |
| кр**о**ты | CR |  | plural | 6 | 5 | 2 | [krɐˈtɨ] |  | moles (burrowing insectivore) |
| леб**е**дь | CR |  | singular | 14.9 | 6 | 2 | [ˈlʲebʲɪtʲ] |  | swan |
| дер**е**во | CR |  | singular | 171 | 6 | 3 | [ˈdʲerʲɪvə] |  | tree, wood |
| вр**а**чи | CR |  | plural | 173 | 5 | 2 | [vrɐˈt͡ɕi] |  | therapists, doctors |
| пл**е**чо | CR |  | singular | 236.4 | 5 | 2 | [plʲɪˈt͡ɕɵ] |  | shoulder, upper arm, brachium, humerus |
| ст**е**на | CR |  | singular | 261 | 5 | 2 | [sʲtʲɪˈna] |  | wall |
| с**е**мья | CR |  | singular | 276 | 5 | 2 | [sʲɪˈmʲja] |  | family |
| л**и**сты | CR |  | plural | 127.6 | 5 | 2 | [lʲɪˈstɨ] |  | sheets (of paper or similar) |
| м**е**ста | CR |  | plural | 926 | 5 | 2 | [ˈmʲestə] |  | places, seats |
| п**а**стух | CR |  | singular | 9.6 | 6 | 2 | [pɐˈstux] |  | shepherd |
| п**е**нёк | CR |  | singular | 9.3 | 5 | 2 | [pʲɪˈnʲɵk] |  | diminutive of пень (penʹ): (small) stump, stub |
| пл**о**ды | CR |  | plural | 47.5 | 5 | 2 | [pɫɐˈdɨ] |  | fruits |
| вр**а**жда | CR |  | singular | 7 | 6 | 2 | [vrɐˈʐda] |  | enmity, hostility, animosity |
| пч**е**ла | CR |  | singular | 10.3 | 5 | 2 | [pt͡ɕɪˈɫa] |  | bee |
| рук**а**ва | CR |  | plural | 42.9 | 6 | 3 | [rʊkɐˈva] |  | sleeves |
| дв**о**рец | CR |  | singular | 60 | 6 | 2 | [dvɐˈrʲet͡s] |  | palace |
| св**и**нья | CR |  | singular | 23.1 | 6 | 2 | [svʲɪˈnʲja] |  | pig, hog, sow, swine |
| син**е**ва | CR |  | singular | 7.9 | 6 | 3 | [sʲɪnʲɪˈva] |  | blue colour, blue |
| ск**а**ла | CR |  | singular | 30.7 | 5 | 2 | [skɐˈɫa] |  | rock, cliff, crag (mass of projecting rock) |
| д**о**бро | CR |  | singular | 59 | 5 | 2 | [dɐˈbro] |  | the good, goods, property |
| сл**о**ны | CR |  | plural | 23.8 | 5 | 2 | [sɫɐˈnɨ] |  | elephants |
| сп**и**на | CR |  | singular | 183.1 | 5 | 2 | [spʲɪˈna] |  | (anatomy) back |
| ст**а**рик | CR |  | singular | 151 | 6 | 2 | [stɐˈrʲik] |  | old man |
| ст**е**кло | CR |  | singular | 102.8 | 6 | 2 | [sʲtʲɪˈkɫo] |  | glass |
| стр**а**на | CR |  | singular | 725.7 | 6 | 2 | [strɐˈna] |  | country |
| т**о**лпа | CR |  | singular | 96.7 | 5 | 2 | [tɐɫˈpa] |  | crowd, throng |
| тр**о**па | CR |  | singular | 16.6 | 5 | 2 | [trɐˈpa] |  | path, trail (a trail for the use of, or worn by, pedestrians) |
| уг**о**лок | CR |  | singular | 32.2 | 6 | 3 | [ʊɡɐˈɫok] |  | diminutive of угол (úgol): small corner (part or region), nook |
| уч**е**ник | CR |  | singular | 80.4 | 6 | 3 | [ʊt͡ɕɪˈnʲik] |  | schoolboy, pupil, apprentice, learner, disciple, follower |
| хв**о**сты | CR |  | plural | 55.4 | 6 | 2 | [xvɐˈstɨ] |  | tails |
| с**и**няк | CR |  | singular | 13 | 5 | 2 | [sʲɪˈnʲak] |  | (pathology) bruise (medical: mark on the skin) |
| в**а**лна | MR | волна | singular | 95.4 | 5 | 2 | [vɐɫˈna] | 2 | wave |
| в**а**рона | MR | ворона | singular | 12.6 | 6 | 3 | [vɐˈronə] | 2 | crow (bird) |
| г**а**лова | MR | голова | singular | 709 | 6 | 3 | [ɡəɫɐˈva] | 2 | head, mind |
| гн**и**здо | MR | гнездо | singular | 20.8 | 6 | 2 | [ɡnʲɪˈzdo] | 3 | nest, aerie, socket, bezel |
| гр**а**за | MR | гроза | singular | 15.7 | 5 | 2 | [ɡrɐˈza] | 3 | thunder, thunderstorm, disaster, danger, menace |
| д**а**жди | MR | дожди | plural | 83.2 | 5 | 2 | [dɐˈʐdʲi] | 2 | rain |
| дв**а**ры | MR | дворы | plural | 166 | 5 | 2 | [dvɐˈrɨ] | 3 | courtyards, homesteads |
| дл**е**на | MR | длина | singular | 67.7 | 5 | 2 | [dlʲɪˈna] | 3 | length |
| зв**а**нок | MR | звонок | singular | 76.5 | 6 | 2 | [zvɐˈnok] | 3 | bell (signaling device such as a bicycle bell), bell (an audible signal such as a school bell), ring, call (a telephone call or telephone conversation) |
| зв**и**зда | MR | звезда | singular | 122 | 6 | 2 | [zvʲɪˈzda] | 3 | star (celestial body), star (celebrity), star (geometric shape) |
| зв**и**рёк | MR | зверёк | singular | 5.3 | 6 | 2 | [zvʲɪˈrʲɵk] | 3 | diminutive of зверь (zverʹ): small beast |
| з**и**мля | MR | земля | singular | 494 | 5 | 2 | [zʲɪˈmlʲa] | 2 | earth, land, ground, soil |
| з**и**рно | MR | зерно | singular | 30.3 | 5 | 2 | [zʲɪrˈno] | 2 | grain, cereal, seed |
| к**а**льцо | MR | кольцо | singular | 59.5 | 6 | 2 | [kɐlʲˈt͡so] | 2 | ring, hoop |
| м**а**зги | MR | мозги | plural | 84.5 | 5 | 2 | [mɐzˈɡʲi] | 2 | brains |
| л**е**сица | MR | лисица | singular | 2.8 | 6 | 3 | [lʲɪˈsʲit͡sə] | 2 | fox (Vulpes vulpes), vixen (female of the fox) |
| л**и**сник | MR | лесник | singular | 3.6 | 6 | 2 | [lʲɪˈsʲnʲik] | 2 | ranger, forest ranger, woodsman |
| с**и**рьга | MR | серьга | singular | 5.7 | 6 | 2 | [sʲɪrʲˈɡa] | 2 | earring |
| м**а**ряк | MR | моряк | singular | 23.2 | 5 | 2 | [mɐˈrʲak] | 2 | seaman, sailor |
| м**а**сты | MR | мосты | plural | 65 | 5 | 2 | [mɐˈstɨ] | 2 | bridges |
| н**а**чник | MR | ночник | singular | 1.6 | 6 | 2 | [nɐt͡ɕˈnʲik] | 2 | nightlight |
| н**и**бeca | MR | небеса | plural | 33 | 6 | 3 | [nʲɪbʲɪˈsa] | 2 | heavens, skies |
| ос**и**нь | MR | осень | singular | 81 | 5 | 2 | [ˈosʲɪnʲ] | 3 | autumn, fall |
| п**е**сьмо | MR | письмо | singular | 304 | 6 | 2 | [pʲɪsʲˈmo] | 2 | letter (written communication), writing (action),  writing system, script |
| п**е**тно | MR | пятно | singular | 50 | 5 | 2 | [pʲɪtˈno] | 2 | spot, blot, blemish, stain |
| м**и**тла | MR | метла | singular | 6.7 | 5 | 2 | [mʲɪˈtɫa] | 2 | broom, besom |
| пов**о**р | MR | повар | singular | 13.3 | 5 | 2 | [ˈpovər] | 4 | cook, chef |
| п**о**руса | MR | паруса | plural | 15.2 | 6 | 3 | [pərʊˈsa] | 2 | sails |
| пт**и**нец | MR | птенец | singular | 4.6 | 6 | 2 | [ptʲɪˈnʲet͡s] | 3 | chick, nestling, fledgling, baby bird |
| с**а**сна | MR | сосна | singular | 30.1 | 5 | 2 | [sɐˈsna] | 2 | pine, pine tree, pine wood |
| р**а**жок | MR | рожок | singular | 5.1 | 5 | 2 | [rɐˈʐok] | 2 | ear trumpet, cone (of ice cream) |
| н**а**здря | MR | ноздря | singular | 12.6 | 6 | 2 | [nɐzˈdrʲa] | 2 | nostril |
| с**и**стра | MR | сестра | singular | 121.3 | 6 | 2 | [sʲɪˈstra] | 2 | sister |
| зл**а**дей | MR | злодей | singular | 10.1 | 6 | 2 | [zɫɐˈdʲeɪ̯] | 3 | malefactor, evildoer, villain, miscreant, scoundrel |
| сл**и**ды | MR | следы | plural | 86.9 | 5 | 2 | [slʲɪˈdɨ] | 3 | track, footprint, footstep, trace, sign |
| ст**а**лы | MR | столы | plural | 402.5 | 5 | 2 | [stɐˈɫɨ] | 3 | tables |
| стр**и**ла | MR | стрела | singular | 20.7 | 6 | 2 | [strʲɪˈɫa] | 4 | arrow, pointer, indicator, (technical): crane arm, jib, overhang beam, boom (of a derrick) |
| тр**о**ва | MR | трава | singular | 88.7 | 5 | 2 | [trɐˈva] | 3 | grass, herb, weed |
| уг**а**лёк | MR | уголёк | singular | 5.5 | 6 | 3 | [ʊɡɐˈlʲɵk] | 3 | diminutive of уголь (úgolʹ): coal, ember (glowing piece of coal or wood) |
| х**а**лмы | MR | холмы | plural | 32.7 | 5 | 2 | [xɐɫˈmɨ] | 2 | hills |
| цв**и**ток | MR | цветок | singular | 92.4 | 6 | 2 | [t͡svʲɪˈtok] | 3 | flower (flowering part of a plant), flower (with stem), flowering plant, potted home plant |
| ч**е**сло | MR | число | singular | 393.5 | 5 | 2 | [t͡ɕɪˈsɫo] | 2 | cardinal number, date, day, (grammar) number |
| ч**и**рвяк | MR | червяк | singular | 4.4 | 6 | 2 | [t͡ɕɪrˈvʲak] | 2 | worm |
| шк**о**фы | MR | шкафы | plural | 48.4 | 5 | 2 | [ʂkɐˈfɨ] | 3 | cupboards, wardrobes, lockers, cabinets  bookcases |
| с**е**лач | MR | силач | singular | 1.5 | 5 | 2 | [sʲɪˈɫat͡ɕ] | 2 | strong man |
| г**а**рбун | MR | горбун | singular | 1.6 | 6 | 2 | [ɡɐrˈbun] | 2 | humpback (humpback person) |
| щ**е**пцы | MR | щипцы | plural | 1.8 | 5 | 2 | [ɕːɪpˈt͡sɨ] | 2 | tongs, pliers, pincers, nippers |
| б**и**гун | MR | бегун | singular | 2.4 | 5 | 2 | [bʲɪˈɡun] | 2 | runner, one who runs |
| л**и**сок | MR | лесок | singular | 2.8 | 5 | 2 | [lʲɪˈsok] | 2 | diminutive of лес (lěs): grove, small wood |
