## Supplementary Information 2. for "Electrophysiological signatures of spelling sensitivity development from primary school age to adulthood"

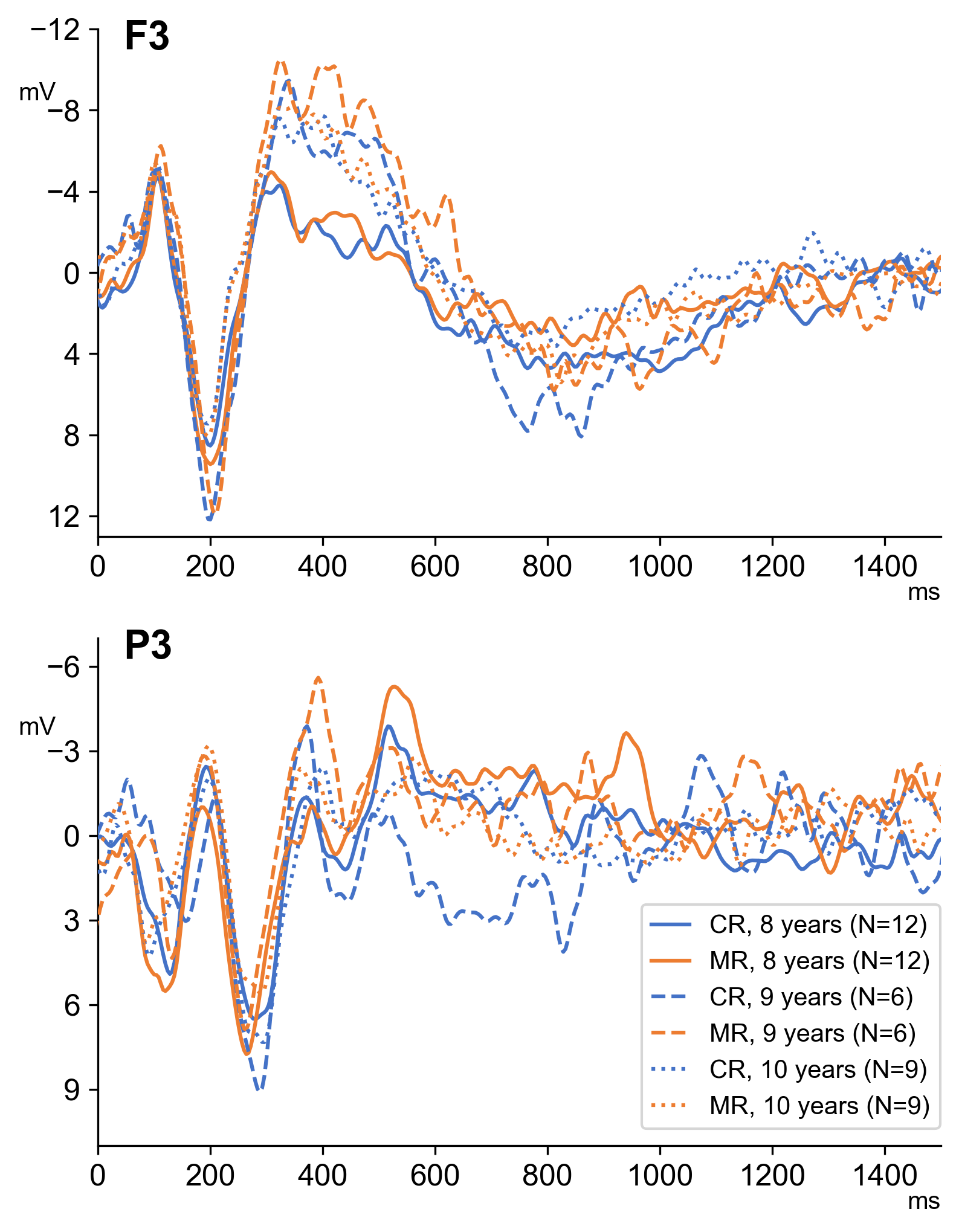

The figure shows the averaged ERPs for correct (CR) and misspelling (MR) words in children aged 8, 9, and 10 for P3 and F3. We did not find any difference between ERPs to CR and MR conditions in these subgroups of children.
